# Divergent coronaviruses discovered in the virome of lamprey with reddening syndrome

**DOI:** 10.64898/2026.04.23.720431

**Authors:** Jessica A Darnley, Stephanie J Waller, Rebecca K French, Riki Parata, Karthiga Kumanan, Luka Finn, Jonah L Yick, Jane Kitson, Patrick Cahill, Ian Davidson, Ryan Hunter, Benjamin W Neuman, Kate S Hutson, Jemma L Geoghegan

**Author notes:** Corresponding author: Jemma L Geoghegan.

## Abstract

Lamprey reddening syndrome (LRS) is an emerging disease affecting pouched lamprey (*Geotria australis*; kanakana/piharau), a culturally and ecologically significant species in Aotearoa New Zealand. Characterised by skin haemorrhaging and elevated mortality, the aetiology of LRS has remained unresolved despite previous investigations. We used a metatranscriptomic approach to characterise viral communities in 28 lamprey from New Zealand and Tasmania, Australia, comparing diseased and presumably healthy individuals. This analysis revealed eight fish-infecting RNA viruses, seven of which were novel, including two highly divergent coronaviruses. One of these coronaviruses possessed a bi-segmented genome structure, and three lamprey were co-infected with both coronaviruses. While these coronaviruses were detected in both healthy and diseased individuals, lamprey with reddening exhibited markedly higher viral abundance, driven by elevated RNA transcripts of both viruses. This pattern suggests that increased coronavirus replication in diseased individuals may be influenced by host stress to environmental factors or co-infection with other pathogens, rather than acting as a sole causative agent of disease. Beyond identifying candidate viral associations, this study expands the known virosphere of an ancient vertebrate lineage and demonstrates the utility of genomics-informed diagnostics for investigating disease in threatened wildlife.

## Introduction

Aotearoa New Zealand’s geographic isolation has generated high levels of endemism across its terrestrial and aquatic biota^1^, yet the microbial communities associated with these species remain largely underexplored^2,3^. Recent work has revealed that endemic wildlife host diverse viromes and microbiomes, many of which appear to exist without overt disease^4–7^. Distinguishing commensal microbial diversity from true or opportunistic disease-causing agents is therefore a central challenge in wildlife disease investigations. This challenge is amplified in threatened species, whose rarity and conservation value make sampling or traditional destructive diagnostic approaches undesirable or untenable.

The pouched lamprey, *Geotria australis*; kanakana or piharau, is a culturally and ecologically significant anadromous fish with deep evolutionary origins^8–10^. In New Zealand, this species has experienced recurring mortality events associated with a condition termed lamprey reddening syndrome (LRS)^11^, characterised by haemorrhagic skin lesions and tissue reddening. Since its recognition in New Zealand in 2011, multiple bacterial taxa have been isolated from affected individuals, yet no single pathogen has been consistently identified as the cause. Similar clinical presentations have been reported in Western Australia^12^, raising questions about shared susceptibility or pathogen transmission between geographically connected populations^13^. The aetiology of LRS remains unresolved despite several investigations over the last 15 years.

Wildlife hosts commonly harbour diverse asymptomatic viral infections. Classical frameworks such as Koch’s postulates^24^ are often insufficient in the context of modern molecular detection, particularly for unculturable or opportunistic pathogens^25–27^. Instead, disease causation can be evaluated through patterns of association, abundance, biological plausibility and ecological context^28^. In aquatic systems, where environmental viral diversity is high and host stressors are typically unknown, disease may arise from interactions among pathogens and the environment, rather than a single pathogenic agent^28–30^.

Traditional diagnostic approaches, including microbial culture and targeted PCR, are inherently limited to detecting known pathogens. In contrast, untargeted genomic approaches such as metagenomics and metatranscriptomics enable comprehensive profiling of the host infectome, including the full complement of viruses present within a sample^14–17^. These methods are particularly powerful for detecting RNA viruses, which evolve rapidly and may evade conventional diagnostics^18–20^. Genomics-informed investigations have transformed disease outbreak analysis in human and animal health settings and are increasingly applied to wildlife disease, where causative agents are often unknown^17,21–23^.

Here, we apply a metatranscriptomic approach to investigate LRS in pouched lamprey sampled from New Zealand and Australia. By comparing diseased and apparently healthy individuals, we aimed to identify viral taxa (i.e. the virome) associated with reddening, assess geographic differences in virome composition and evaluate whether LRS is consistent with viral infection. In doing so, we demonstrate the utility of genomics-informed diagnostics for resolving complex disease syndromes in culturally significant and evolutionarily distinct wildlife species.

## Methods

### Animal ethics

This project was approved by the Nelson Marlborough Institute Technology Animal Ethics Committee (AEC permit AEX-2023-CAW-08). This project was co-designed from its inception by members of the Hokonui Rūnanga, who represent the indigenous peoples of the area from where animals were sampled in New Zealand. Samples of *Geotria australis* in Australia were provided by the Inland Fisheries Service (IFS), Tasmania, working under the *Inland Fisheries Act 1995*.

### Kanakana sampling

Pouched lamprey (kanakana or *Geotria australis*) were collected from the Te Au-nui pihapiha-kanakana falls, Mataura River (-46.19026, 168.87141), in the South Island of New Zealand between 5 September and 29 October 2024. Lamprey were hand gathered by members of Hokonui Rūnanga, indigenous people of the area, wearing cotton gloves, continuing the traditional harvest method by early indigenous peoples. Lamprey collected included presumably healthy animals (n = 2), animals with very slight reddening but otherwise healthy (n = 5), and animals exhibiting external skin abnormalities consistent with LRS (n = 10) (Figure 1). Lamprey were first anaesthetised with a dose of 50 ppm AQUI-S then killed using an overdose of AQUI-S (at least 100 ppm). To avoid cross-contamination, a separate sterile blade was used to excise a piece of tissue approximately 3mm^3^ from the following organs: skin, gill, kidney, liver and muscle tissue (Supplementary Table 1). Tissue was stored in separate, labelled 2-mL tubes containing RNALater (Thermo Fisher Scientific) prior to being couriered on ice to the University of Otago and stored at -80 degrees Celsius.

**Figure 1.**
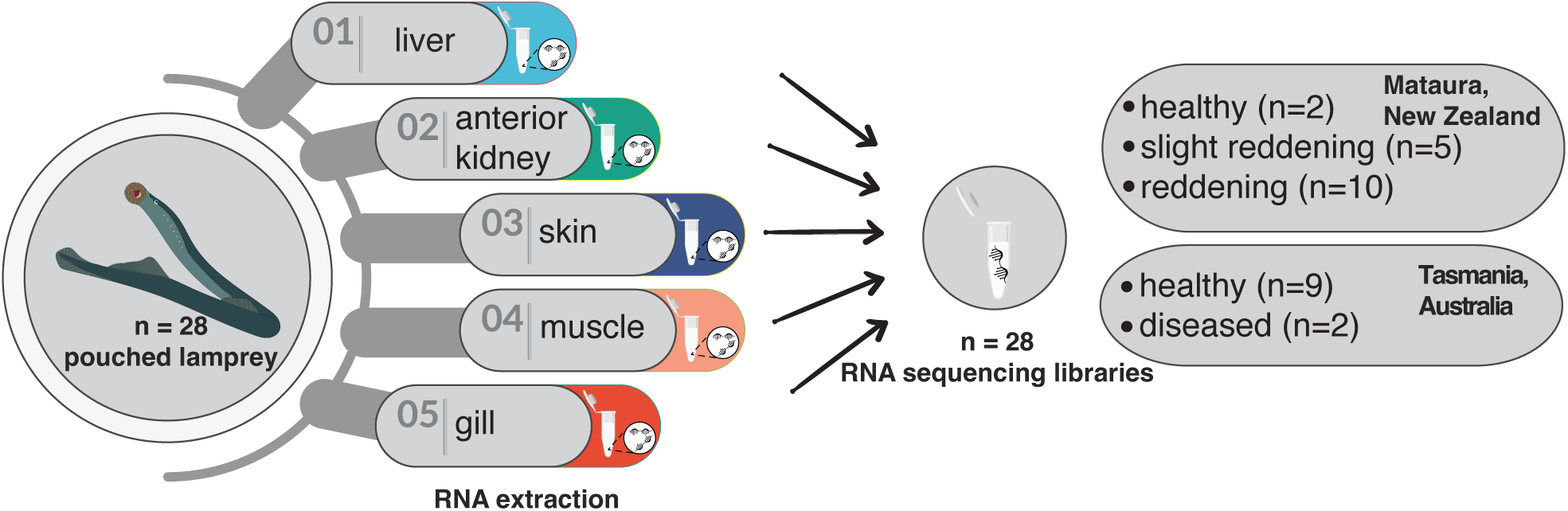
Sampling and RNA pooling protocol. Five tissue samples were homogenised during RNA isolation into a single sample, resulting in a total of 28 sequencing libraries.

In Australia, lamprey were sampled on 4 December 2024. Lamprey were collected from a fish trap maintained by the IFS, installed at the base of the Meadowbank Dam at the head of the River Derwent, Tasmania (-46.61002, 146.84501). Lamprey (n = 11) were euthanised by severing the notochord before being placed on a clean tray. Aseptic techniques were used to obtain organ samples as described for sampling in New Zealand. Nine lamprey were presumably healthy, while two individuals exhibited minor skin abnormalities, including one individual that exhibited haemorrhage around the branchiopores.

### Tissue total RNA extraction

Total RNA was extracted from tissue using the Qiagen RNeasy Plus Mini Kit (Qiagen). Pieces of defrosted tissue were cut using a sterile scalpel and forceps and placed into the lysis buffer. Next, 2-Mercaptoethanol was added to buffer RLT+ at a concentration of 1% and reagent DX was added at a concentration of 0.05% during the tissue lysis stage. A Qiagen TissueRuptor with sterilised probes was used to homogenise each sample for ∼60 seconds. Any tissue debris was removed via centrifugation and the rest of the protocol was followed according to the manufacturer’s directions. RNA was measured using a NanoDrop spectrophotometer (Thermo Fisher) to gauge concentration and purity.

### RNA pooling and sequencing

Extracted RNA was pooled by individual using standard volumes of 15 µL. In cases of duplicated tissue, the one with higher RNA concentration was used. The pooled concentration was re-measured (see Supplementary Table 1 and Figure 1). Pooled RNA was subject to RNA sequencing, using the Illumina Stranded Total RNA Prep with Ribo-Zero plus kit (Illumina) for library preparation. Libraries were sequenced on the Illumina NovaSeqX platform and 150 bp paired-end reads were generated.

### Virus assembly and discovery

Raw sequencing data were trimmed using Trimmomatic (v0.38)^31^, to remove the sequencing adaptors. Library quality was checked using FastQC (v0.12.0)^32^, before *de novo* assembly using MEGAHIT (v1.2.9)^33^. Protein-level homology searches were used to annotate the assembled contigs against NCBIs non-redundant protein database using DIAMOND BLASTx (v2.1.9) (Basic Local Alignment Search Tool).

Potential viral hits were manually screened to identify fish host-infecting viruses via manual inspection and translation of open reading frames (ORFs) in Geneious Prime (v2025.0.3) (https://www.geneious.com), and additional BLASTn and BLASTp searches (https://blast.ncbi.nlm.nih.gov/Blast.cgi) were used to confirm virus transcripts and remove false positive hits. If a contig’s closest match e-value was <1×10^-5^ and the top blast hits were a vertebrate-infecting virus, then the contig was considered a putative lamprey virus.

### Estimating viral transcript abundance

Abundance estimations were made by mapping sequence reads using Bowtie2^34^ to the assembled contigs. Reads per million (RPM) were calculated by multiplying standardised library read counts by one million. Read depth and abundances were explored using R (v4.4.2). Shannon diversity and richness were explored using the vegan (v 2.7.1)^35^ R package, and tested with a Kruskal-Wallis test and a post-hoc Dunn test with Benjamini-Hochberg (BH) correction using FSA (v0.10.0)^36^. Non-metric Multidimensional Scaling (NMDS) plots with Bray-Curtis distances were used to visualise community composition measured with Beta diversity and tested using pairwise PERMANOVA with pairwise Adonis (v0.4.1). Plots were generated using ggplot2 (v3.5.2)^37^. All R code for statistical analysis is available on GitHub (github.com/Jess-AD/Lamprey).

### Virus phylogenetic analysis

NCBI Taxonomy (https://www.ncbi.nlm.nih.gov/taxonomy) was used to obtain a range of known virus species for each respective taxonomic family or order. Geneious Prime (v2023.2.1) (https://www.geneious.com/) was used to align the amino acid sequences using the MAFFT^38^ (v1.5.0) (104) plugin with the L-INS-i algorithm used for all sequences to ensure the alignment of conserved sequence motifs. TrimAl^39^ (v1.2) was used to trim the alignments and remove ambiguous sequences. IQ-TREE^40^ (v1.6.12) was used to generate maximum likelihood phylogenetic trees using an LG model and FigTree (v1.4.4) (http://tree.bio.ed.ac.uk/software/figtree/) was used to annotate them, which were rooted by the midpoint for clarity only. Figures were annotated further using Adobe Illustrator (v28.1).

Viruses were considered novel species if the highly conserved DNA polymerase or RNA-dependent RNA polymerase (RdRp) region shared <90% amino acid similarity with the closest related virus in the NCBI database.

### Assembly and phylogenetic analysis of full genomes from the Coronaviridae family

Full genomes were assembled by *de novo* assembly of overlapping reads. Once a consensus sequence was assembled, other genomes were assembled by mapping reads back to the consensus. To annotate full genome sequences for the two coronaviruses, ORFs were translated and subject to HHpred^41^ analysis, or BLASTp searches against a curated database of known viral structural and accessory proteins. Motifs for each protein were identified at the protein sequence level.

To account for the possibility of different evolutionary rates within the *Coronaviridae* replicase polyprotein leading to misleading phylogenies, the phylogenetic tree was assembled using only the N-terminal nidovirus-specific extension (NiRAN), RdRp, zinc binding domains (ZBD) and Helicase (Hel) regions, which are known to be conserved at the same rate^42^, providing a reliable phylogeny. Due to the high divergence of the two identified coronaviruses, a consensus sequence of each virus was used to re-map sequence reads using Bowtie2^34^ to determine the abundance of each coronavirus in each library. To account for misassignment, reads were standardised to RPM and subject to an abundance cut-off of one RPM. In addition, to view the evolution of the coronaviruses, maximum likelihood phylogenetic trees were estimated using nucleotide alignments of each species using an HKY model in IQ-Tree^40^.

### Novel virus nomenclature

Novel viruses were named after discussion with representatives of the Hokonui Rūnanga and InlandFisheries Tasmania, where the fish with this virus were collected. *Geotria australis* was chosen for the prefix as the viruses found were shared between New Zealand and Australia, and likely to be found shared in future investigations; therefore, a non-specific location name was chosen.

## Results

### Classification of Lamprey Reddening Syndrome

Lamprey collected from Te Au-nui pihapiha-kanakana falls, Mataura River, Mataurain the South Island of New Zealand were classified as exhibiting reddening syndrome if they showed bleeding, bruising or reddening of the skin (n = 10) (Figure 2a-d). Five fish (n = 5) were considered healthy yet ‘compromised’ with only slight reddening, while two were presumably healthy with no evidence of skin disease (n = 2) (not pictured). Fish in Tasmania were presumably healthy (n = 9), but two (n = 2) fish showed external signs of an unknown, potentially related, disease and were considered ‘diseased (Figure 2e-f).

**Figure 2.**
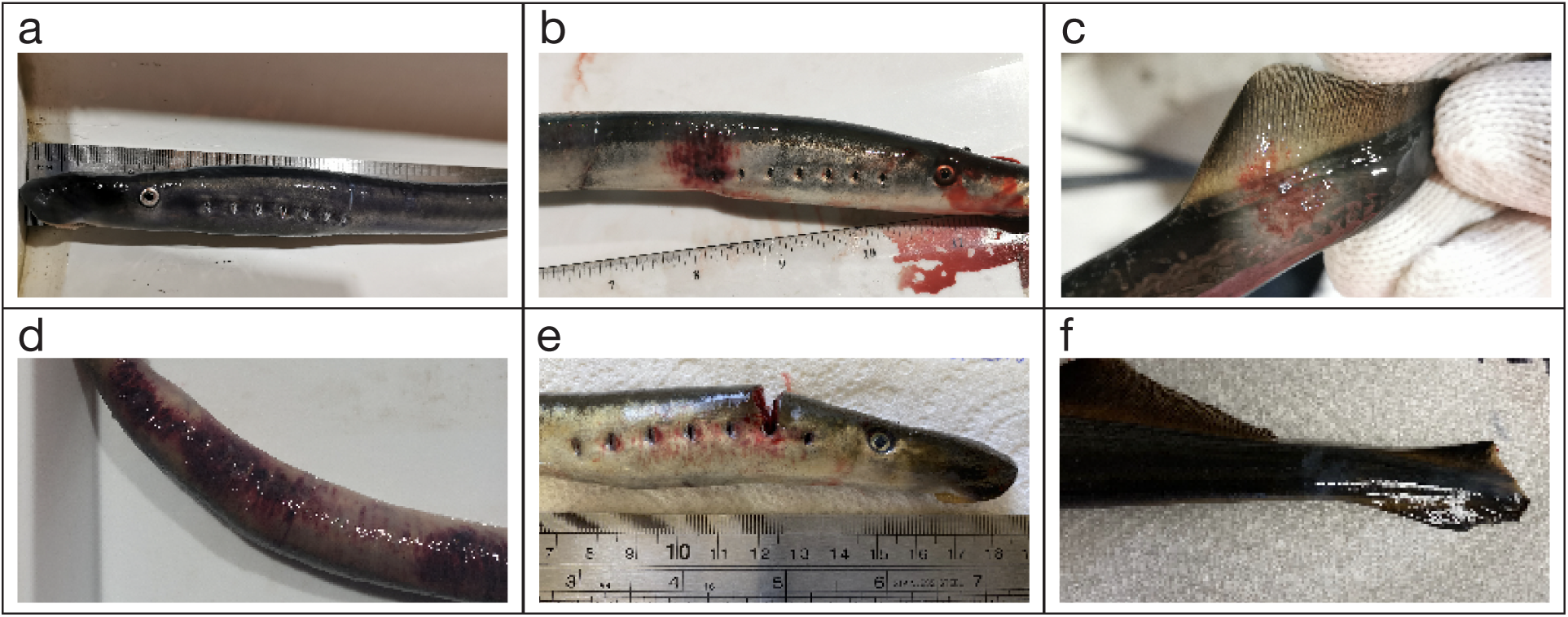
Photographs of sampled lamprey representative of disease states: healthy (**a**); exhibiting reddening syndrome (**b, c, d**) and compromised with only slight reddening (**e**). Disease status and gross evidence of disease included: (**a**) healthy with no evidence of reddening, (**b**) abdominal reddening, (**c**) d reddening at the base of the tail fin and (**d**) widespread reddening. Lamprey sampled from Tasmania considered compromised (**e**) exhibited red spots around the branchiopores (note severed notochordis associated with euthanasia of the animal) and (**f**) a fungal lesion on skin near the tail fin. Photographs courtesy of Hokonui Rūnanga and the Cawthron Institute.

Between three and five tissues were biopsied from each fish. Total RNA was extracted from each tissue type and pooled per individual, comprising a median pooled RNA concentration of 164.3 ng/µL. Total RNA sequencing yielded, on average, 101 million reads per library (Figure 3).

**Figure 3.**
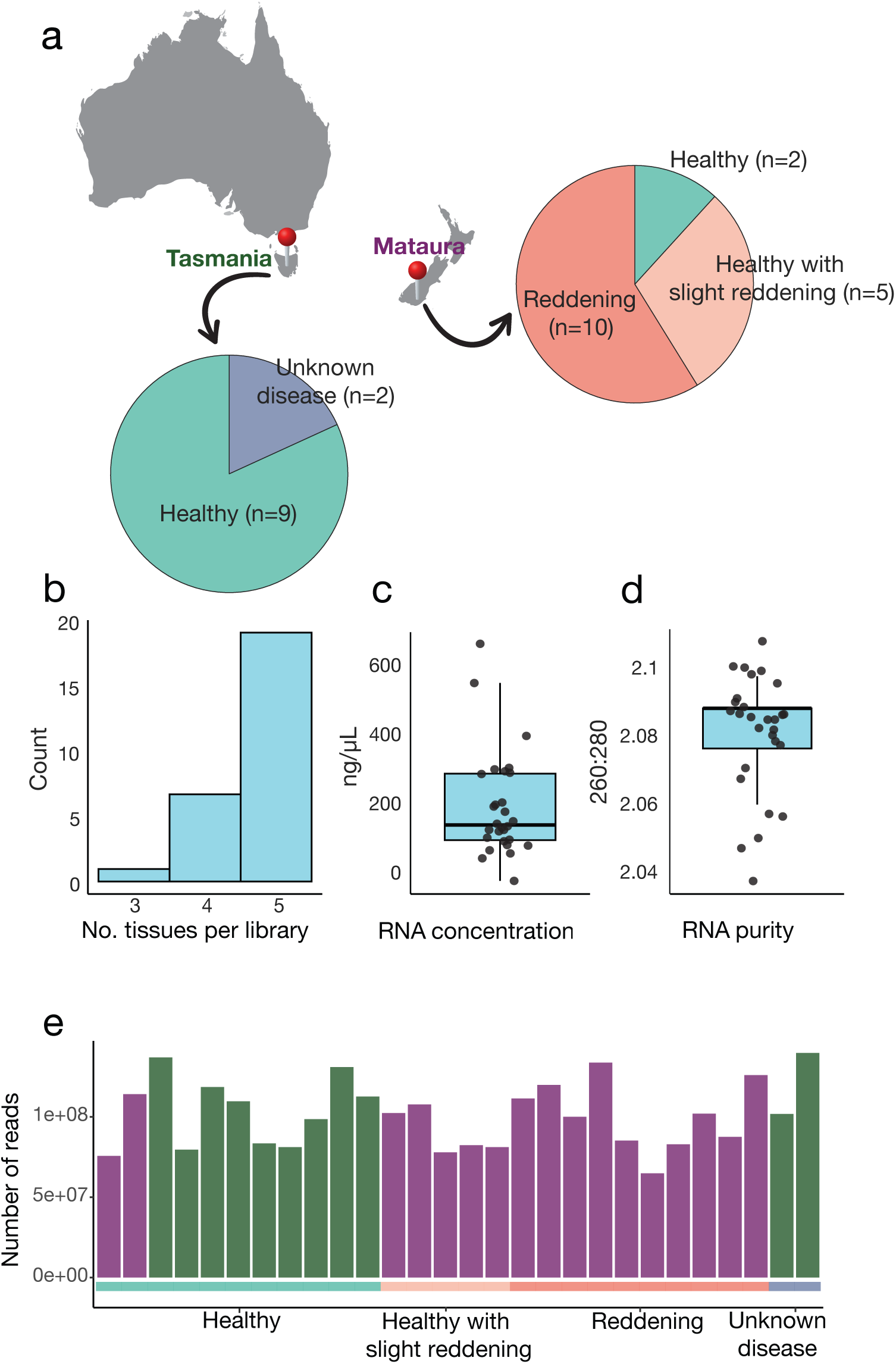
Sampling, RNA extraction and sequencing results. (**a**) Map of Australia and New Zealand showing the two sampling locations in Tasmania, Australia and Mataura, New Zealand with the number of individuals of each health status indicated within the pie charts. (**b**) Bar graph showing the number of tissues per individual in each library. (**c**) Boxplot of RNA concentration for each pool with raw data superimposed. (**d**) Boxplot of RNA purity (260:280 ratio) for each pool with raw data superimposed. Boxplots show the median (thick horizontal line), interquartile range (box), whiskers extending to the most extreme data points within 1.5x interquartile range and individual points are plotted. (**e**) Bar graph showing the number of reads sequenced per library, with samples from Tasmania (green) and Mataura (purple).

### Diversity and abundance of kanakana viruses

Using a metatranscriptomic approach, we revealed eight fish-infecting viral species, all of which were RNA viruses and seven were novel species meaning that they shared less than 90% amino acid sequence similarity within their most conserved gene (i.e. RNA-dependent RNA polymerase, RdRp). Additionally, we uncovered the full genome of a novel virus that likely infects invertebrate hosts, which may be associated with eukaryotic parasites, the environment or lamprey diet. Viruses from the *Coronaviridae* family were found at high relative abundances across multiple sequencing libraries (Figure 4). While viruses from the *Astroviridae* were also found across multiple libraries, these were identified at much lower relative abundances, and viruses from the *Orthomyxoviridae* and the *Picornaviridae* were found in one library only.

**Figure 4.**
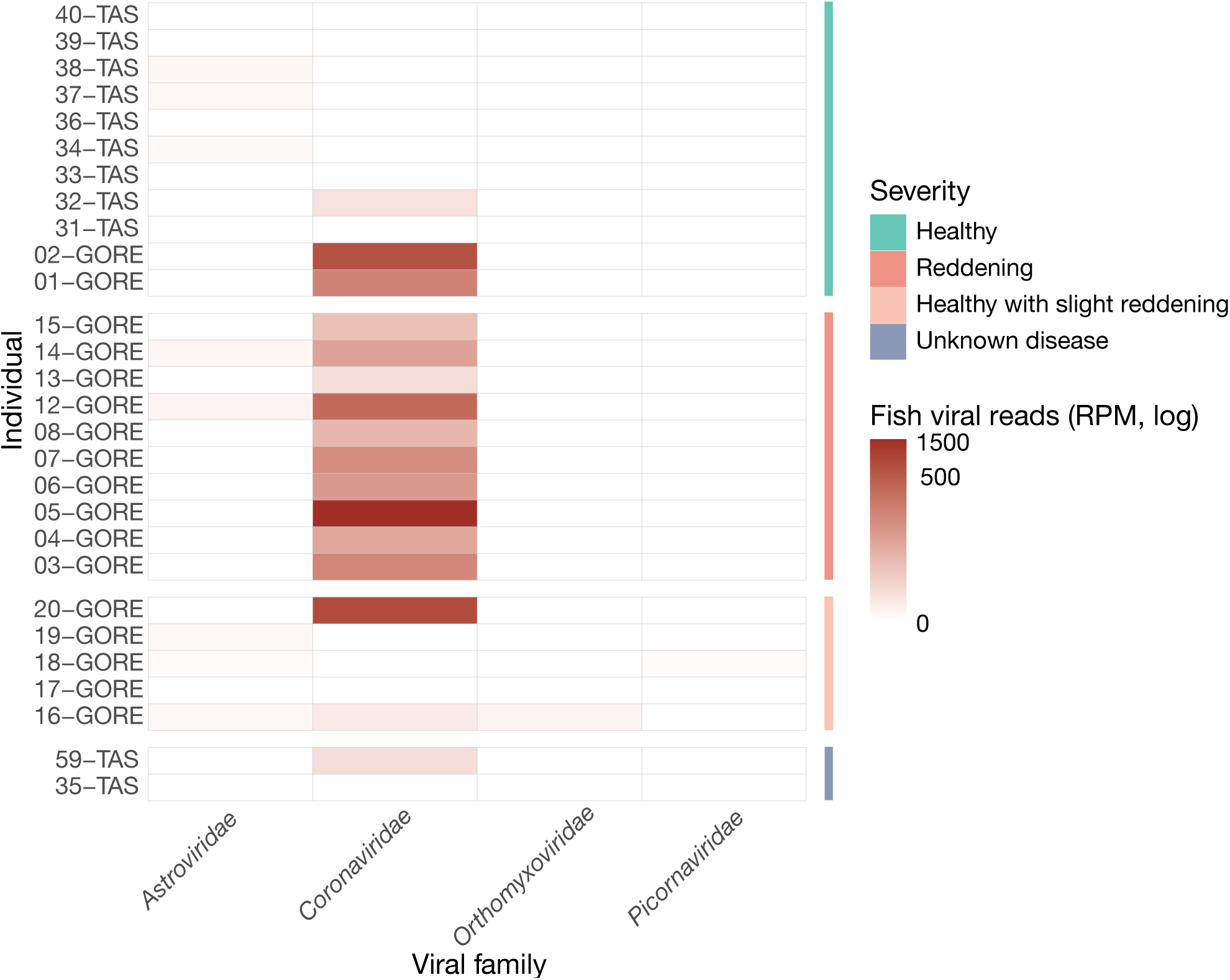
Viral reads per million (RPM, log) for fish-infecting viruses detected using metatranscriptomics, grouped by health status. Each row represents an individual fish, and each column a viral family. RPM values are indicated by colour intensity, with white = no reads, dark red = high abundance.

Standardised abundances of fish-infecting viruses differed significantly among health status categories (Kruskal–Wallis test, χ² = 9.75, *p* = 0.021; Figure 5). To test which groups contributed to this difference, a post-hoc Dunn’s test revealed that fish with LRS (or reddening) had significantly higher viral abundances compared with healthy fish (BH adjusted *p* = 0.025), while no other pairwise comparisons were statistically significant.

**Figure 5.**
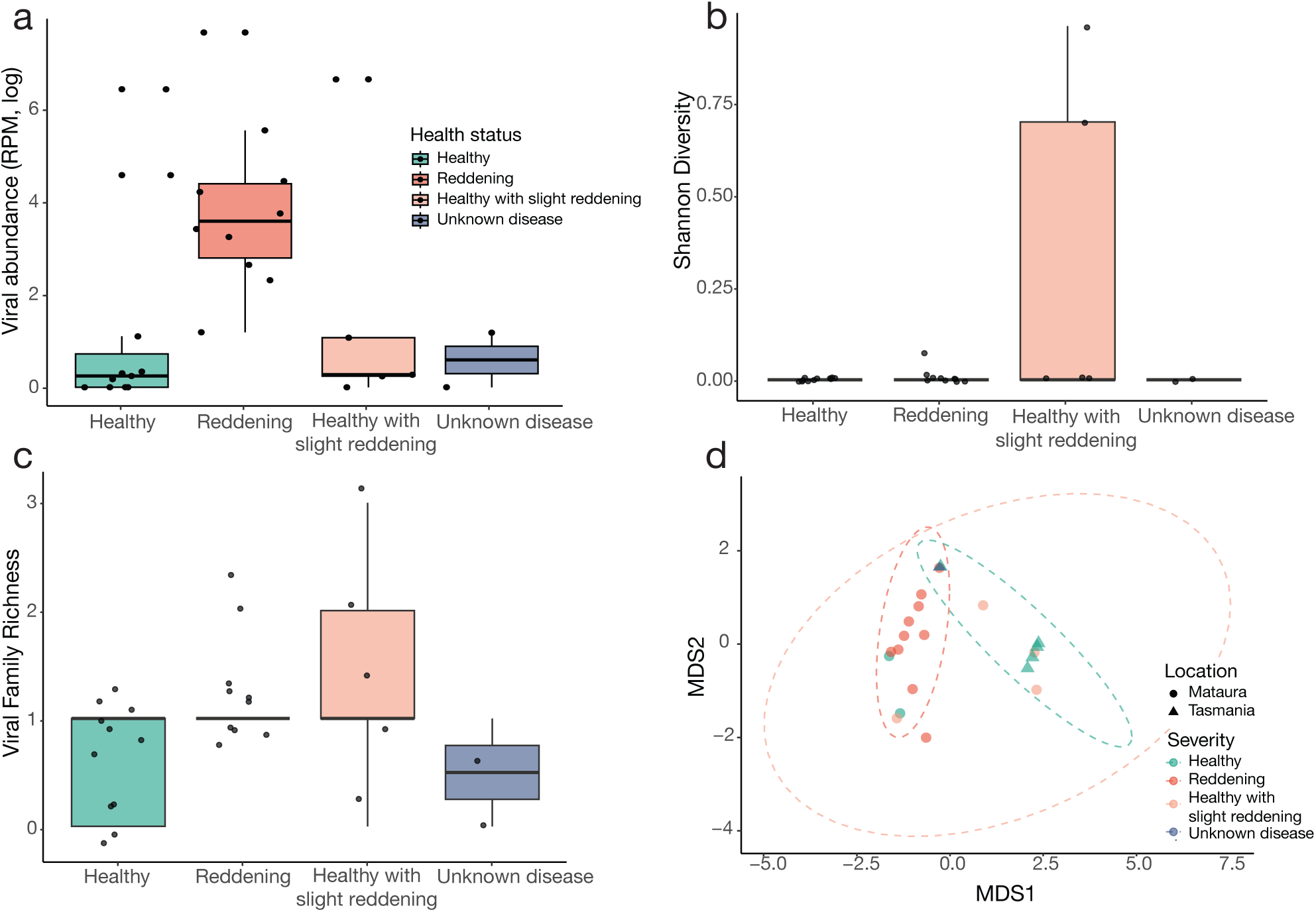
Statistics of virome analysis. (**a**) Logged abundance of viral families by health status. Boxplots show the median (thick horizontal line), interquartile range (box), whiskers extending to the most extreme data points within 1.5x interquartile range and individual points are plotted. Viral abundance differs between health status groups (p = 0.021). (**b**) Shannon diversity of viral communities. (**c**) Richness of viral families for each health status. (**d**) Non-metric Multidimensional Scaling (NMDS) ordination of viral communities across health status and locations. Points represent individual libraries, coloured by health status. PERMANOVA indicated significant differences in community composition (R² = 0.41, p = 0.001), while dispersion tests confirmed that group spread was not significantly different (p = 0.422).

Alpha diversity, measured here using the Shannon index, quantifies both the richness and evenness of viral taxa within each sample, providing an estimate of virome complexity. We found that alpha diversity did not significantly differ among fish with different health statuses (Figure 5). Kruskal-Wallis tests indicated no significant differences in Shannon diversity (χ² = 5.41, *p* = 0.14) or viral richness (χ² = 6.57, *p* = 0.08) across the four classes of fish health status. A post-hoc Dunn test with BH correction confirmed that no pairwise comparisons were statistically significant, indicating that overall viral diversity and viral richness were broadly consistent regardless of reddening.

In contrast, viral community composition, measured via beta diversity, differed significantly across fish health status and sampling location (Pairwise Permutational Multivariate Analysis of Variance, PERMANOVA *p* = 0.003; Figure 5). Beta diversity based on the Bray-Curtis dissimilarity matrix measures how different the composition of viral communities is between samples, accounting for both the identity and relative abundances of viral taxa. Analysis of multivariate dispersion (betadisper) indicated no statistically significant differences in within-group variability (*p* = 0.42), suggesting that the observed differences are due to shifts in community composition rather than unequal spread among groups. PERMANOVA analysis revealed specific group differences, whereby viral communities differed between healthy and reddening (*p* = 0.003) and between reddening and healthy with slight reddening (*p* = 0.005). For sampling locations, lamprey in Tasmania were also significantly different (*p* = 0.005). These results indicate that both health status and location significantly shaped viral community composition in the fish sampled, with pairwise tests highlighting that fish with reddening are the drivers of the overall differences.

### Phylogenetic analysis of kanakana viruses

We next undertook a phylogenetic analysis of the novel lamprey-infecting viruses to place them within their evolutionary contexts (see Supplementary Table 2). Using the most conserved genetic region, the RdRp, we estimated phylogenetic trees for each viral family identified here. We identified four novel viruses within the *Astroviridae* family that appear closely related to other fish-infecting viral species (Figure 6). Geotria australis piscistrovirus 1 was detected in healthy Tasmanian fish, with an average abundance of 0.22 RPM across the two libraries in which it was present. The closest BLASTp hits for the RdRp of this virus was different for the two libraries, with a low amino acid identity of 31% with *Jingmen bat astrovirus 1* (WOK58275.1) and 29% with *Wenling gobies fish astro-like virus* (AVM87598.1) (113). Despite these differences, these two segments shared more than 90% amino acid similarity of their RdRp regions and are considered the same virus.

**Figure 6.**
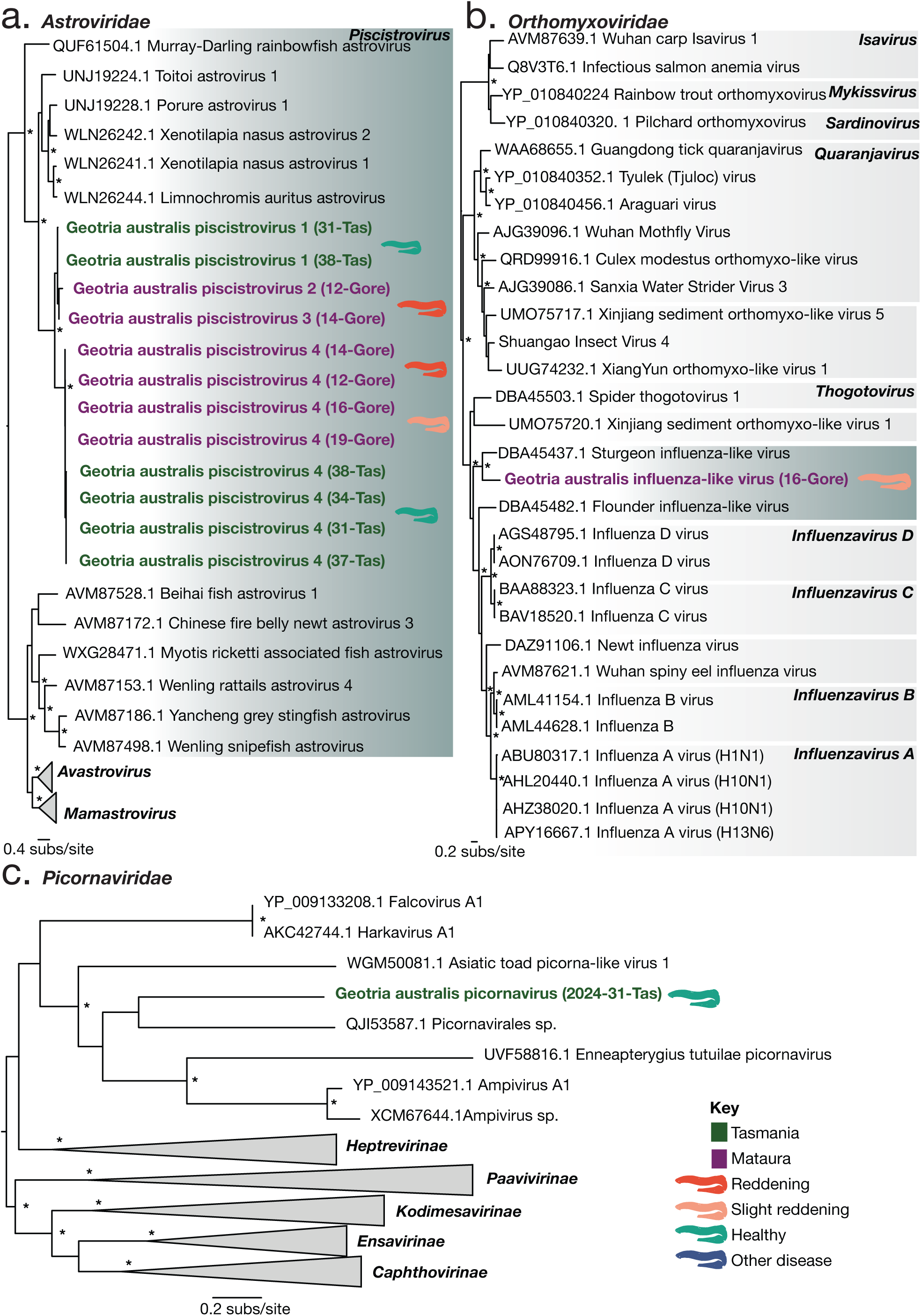
Maximum likelihood phylogenetic trees of RdRp from fish-infecting viruses revealed via metatranscriptomic sequencing. An asterisk (*) indicates ultra-fast bootstrap values of ≥95%. Viruses revealed in the lamprey tissue are bold and coloured green (Tasmanian origin) and purple (Mataura origin). Phylogenetic trees represent (**a**) *Astroviridae*, (**b**) *Orthomyxoviridae* and (**c**) *Picornaviridae*. Some clades have been collapsed to genus or family level, and trees are mid-point rooted for clarity only. Branch lengths are scaled to the number of amino acid substitutions per site. Viruses from individuals with different health statuses are indicated by the colour key.

One lamprey with reddening carried a viral segment that matched closest with 46% amino acid identity to *Jingmen bat astrovirus 1* (WOK58275.1), a virus found in bats with a potential for host jumping (114). This virus was named Geotria australis piscistrovirus 2. Geotria australis piscistrovirus 3 was found in one library from a fish with reddening from Mataura. It was present at 0.24 RPM with a 46% amino acid identity with *Jingmen bat astrovirus 1* (WOK58275.1).

Geotria australis piscistrovirus 4 was found at a combined 1.54 RPM in multiple libraries from Tasmania and Mataura, and found in healthy fish, those with an unknown disease from Tasmania, and healthy fish with slight reddening from Mataura. Piscistrovirus 4 has multiple hits to BLASTp searches with each library’s segment, with 31% similarity to *porure astrovirus* (UNJ19229.1), 40% to *porure astrovirus* (UNJ19228.1), 27-30% to *Xenotilapia nasus astrovirus 2* (WLN26242.1), 39-43% to *Limnochromis auritus astrovirus* (WLN26244.1). All these related viruses infect fish, likely not causing overt disease, with two found in fish from East Africa and one from New Zealand. Differences in detection for the same novel viruses illustrate both their high divergence from known viruses and the substantial number of viruses that remain undiscovered.

One novel virus from the *Orthomyxoviridae* was identified from a fish that was healthy with slight reddening from Mataura (Figure 6) and fell within a clade of other fish-infecting viruses. This virus was found at 0.52 RPM and was tentatively named Geotria australis influenza-like virus after its closest BLASTp hit of 36% similarity, Sturgeon influenza-like virus (DBA45437.1), a virus that has unknown disease associations^43^.

A novel virus, tentatively named Geotria australis picornavirus was detected from the *Picornaviridae* family, falling within an unclassified genus (Figure 6). This virus was found in a healthy Tasmanian lamprey at an abundance of 1.6 RPM. Whilst its closest relative is a virus from a bird swab (QJI53587.1) with 32% similarity, upon phylogenetic analysis, the next closest relatives are aquatic vertebrate hosts. This virus is unlikely to cause disease as it was found in an apparently healthy individual at low abundance.

Two viruses were identified from the *Coronaviridae*, falling within clades of other fish-infecting viruses (Figure 7). Geotria australis coronavirus is a novel virus found within samples collected from Mataura (with an average abundance of 264 RPM) as well as one Tasmanian (with an abundance of 2 RPM) fish with an unknown, but possibly LRS-like, disease. Geotria australis coronavirus is highly divergent from other known coronaviruses. When the consensus sequence is subject to a BLASTp search, it has only 34% amino acid sequence similarity to its closest known genetic relative, Infectious bronchitis virus (IBV, XKR89371.1) and may be suggestive of a new subfamily within the *Coronaviridae*. IBV is a respiratory disease-causing virus in chickens^44^, however upon phylogenetic analysis, the virus fell closer to other fish viruses. We have named this virus Geotria australis coronavirus following discussion with indigenous guardians of this species.

**Figure 7.**
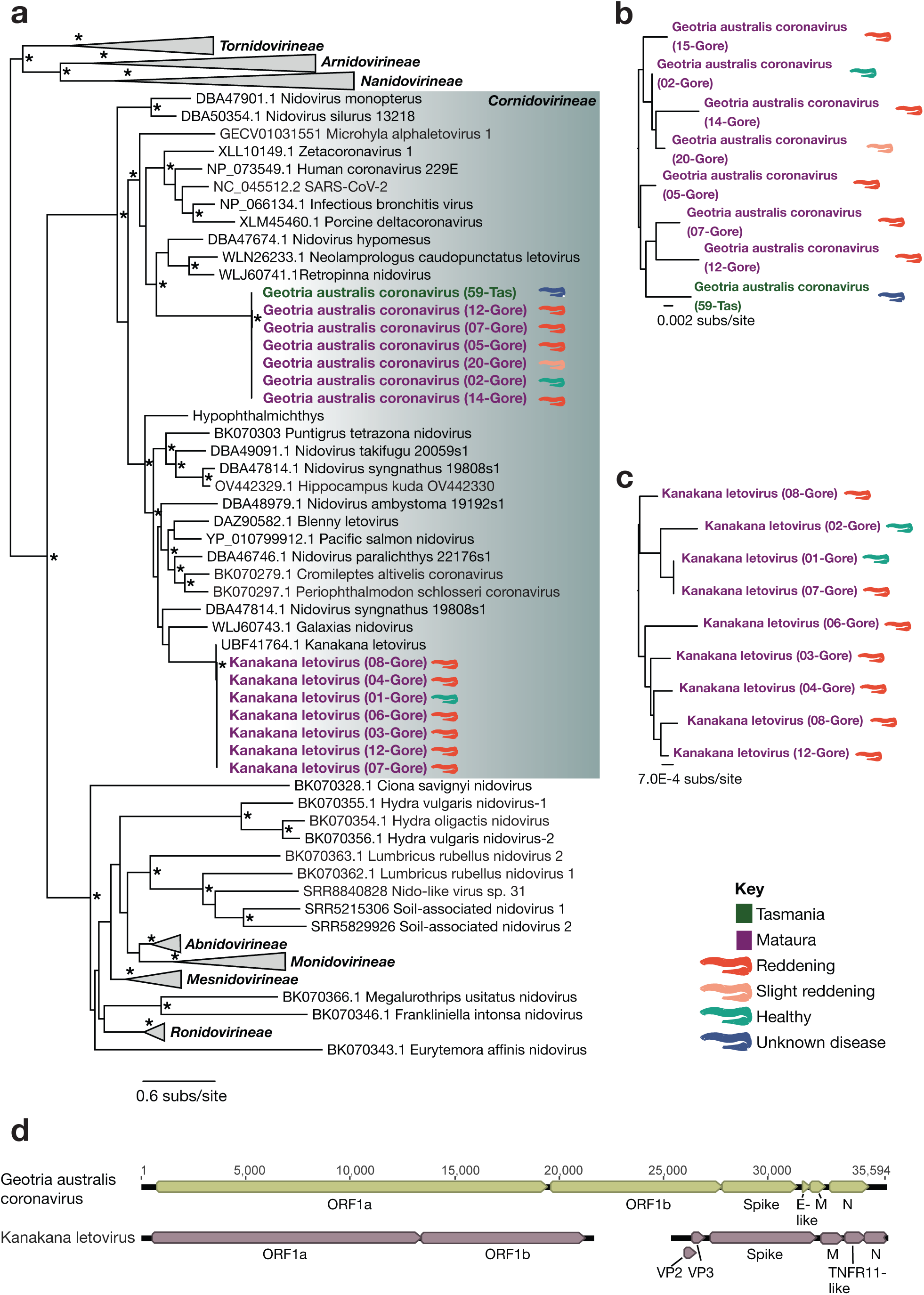
(**a**) Maximum likelihood phylogenetic tree of the Nidovirales order from the NiRAN, RdRp, ZBD and Hel regions with viruses revealed via metatranscriptomic sequencing in bold. Viruses within the shaded box fall within the Cornidovirineae sub-order, and therefore the Coronaviridae family. An asterisk (*) indicates ultra-fast bootstrap values of ≥95%. Note only viruses with complete RdRP regions are shown in the tree. Some clades have been collapsed, and the tree is mid-point rooted for clarity only. (**b**) Whole-genome nucleotide-level phylogenetic tree of Geotria australis coronavirus virus. (**c**) Whole-genome nucleotide-level phylogenetic tree of Kanakana letovirus. Branch lengths are scaled to the number of (**a**) amino acid or (**b-c**) nucleotide substitutions per site. Viruses from individuals with different health statuses are indicated by the colour key. (**d**) Genome structures of the two coronaviruses identified in this study, annotated using sequence and protein homology searches.

*Kanakana letovirus* was present in seven reddening and two healthy fish from Mataura with a mean abundance of 35.8 RPM in fish with reddening, and 8.5 RPM in healthy fish. *Kanakana letovirus* was initially discovered in New Zealand lamprey in 2021 during an initial screening of individuals with LRS on historical samples from 2011^45^; however the recovery of full genomes in the current study has revealed this virus has a bi-segmented genome (Figure 7), a feature only discovered recently in the *Pitovirinae* subfamily^46^. ORF1ab, containing the RdRp, were present on one segment, and the structural and accessory proteins on the second segment. These segments are packaged into two separate envelopes, with infection from both viral particles required for viral replication.

In contrast, Geotria australis coronavirus has a standard coronavirus genome structure, with a monopartite positive-sense single-stranded genome containing ORF1ab and all structural and accessory proteins. It is unusual in size, however, at 35,594 nucleotides in the consensus sequence, with vertebrate-infecting coronaviruses generally possessing a genome size of 26-32 kb^47–48^. Both *Kanakana letovirus* and Geotria australis coronavirus are highly divergent from each other and suggest two separate viral host jumps from other fish species into lamprey. From the eight full genomes assembled of Geotria australis coronavirus, and the nine full genomes assembled of *Kanakana letovirus,* nucleotide-level phylogenetic trees show the high level of divergence between each viral species (Figure 7). Co-infection with both viruses was detected in three lamprey (two with reddening and one healthy).

### Exploration of the coronaviruses

Since the coronaviruses were highly abundant and divergent, we explored their association with health status more thoroughly. We first mapped sequencing reads to the full coronavirus consensus genomes obtained to determine the abundance of each virus in each library (Figure 8). We found that the abundance of *Kanakana letovirus* differed significantly among fish health status (Kruskal–Wallis χ² = 10.30, *p* = 0.016), with a Post-hoc Dunn’s tests with BH correction indicating that reddening fish had higher abundance of *Kanakana letovirus* than healthy fish (adjusted *p* = 0.038) and higher than those with slight reddening (adjusted *p* = 0.048). In contrast, the abundance of Geotria australis coronavirus did not differ significantly among health status (Kruskal–Wallis χ² = 3.2794, *p* = 0.076), and no other pairwise Dunn’s comparisons were statistically significant (all adjusted *p* > 0.05). These results suggest that *Kanakana letovirus* abundance may be associated with health status, while Geotria australis coronavirus has only a trend but no true association.

**Figure 8.**
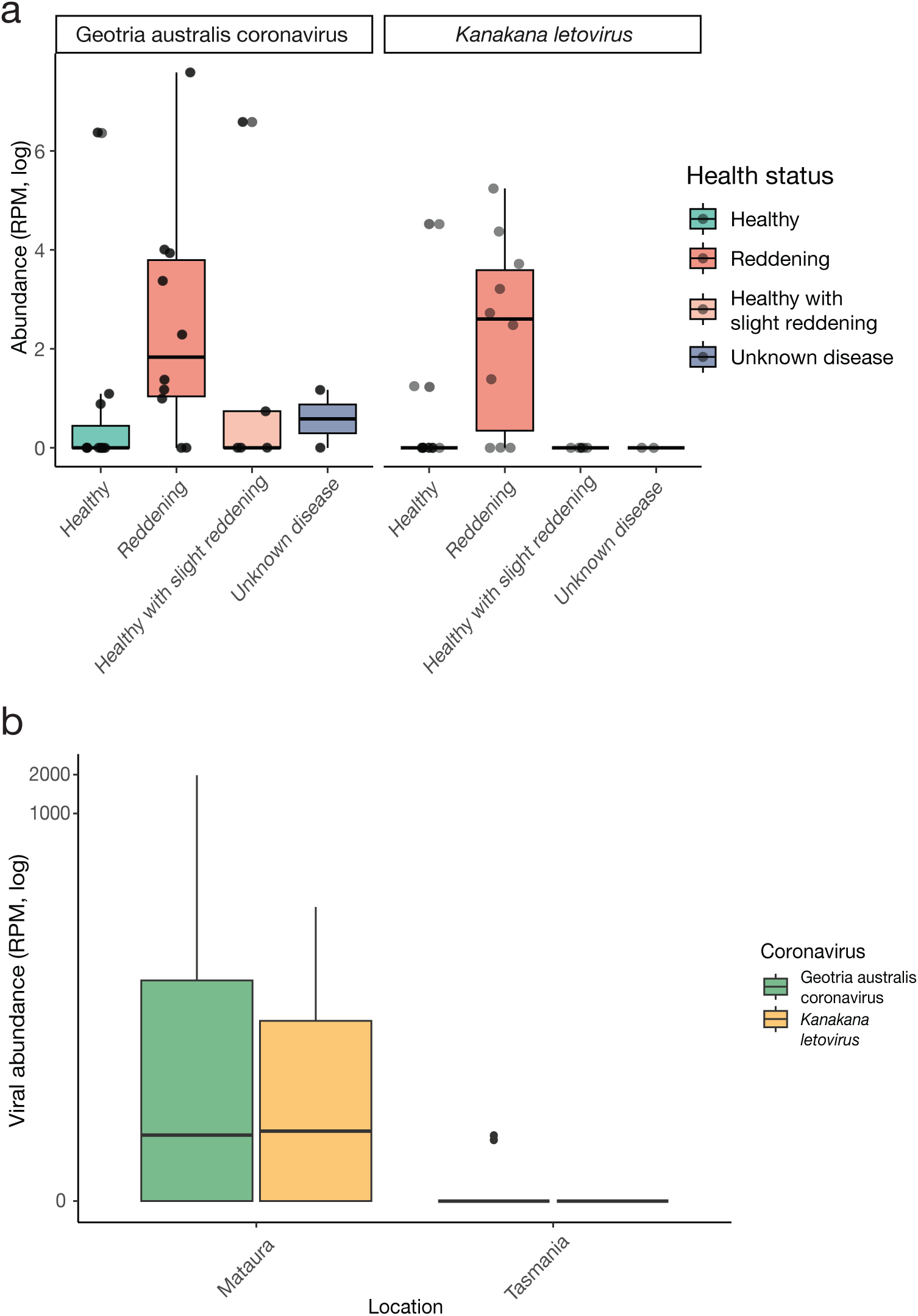
(**a**) Logged abundances (RPM) of *Kanakana letovirus* and Geotria australis coronavirus in each health status group. Boxplots show the median (thick horizontal line), interquartile range (box), whiskers extending to the most extreme data points within 1.5x interquartile range and individual points are plotted. (**b**) logged abundance of *Kanakana letovirus* and geotria australis coronavirus by sampling location.

To determine if the reddening and slight reddening had a similar viral association with disease, the two were combined, and abundance comparisons made between the new, combined group with healthy and unknown diseased. The abundance of *Kanakana letovirus* was not significant with this new health status grouping (Kruskal–Wallis χ² = 6.88, *p* = 0.194), and neither was the abundance of Geotria australis coronavirus (Kruskal–Wallis χ² = 4.4322, *p* = 0.109). Combining the abundance of both coronaviruses with health status showed borderline association with disease (Kruskal–Wallis χ² = 5.9892, *p* = 0.05006), but no true statistical significance. This suggests a trend between coronavirus infection and LRS, but a coronavirus infection is likely not the only pathogen involved with this disease.

When comparing coronavirus abundance with sampling location (Figure 8), viral abundance differed significantly. Both *Kanakana letovirus* (Wilcox test, *p* = 0.005) and Geotria australis coronavirus (Wilcox test, *p* = 0.006) showed higher abundance in Matura compared with Tasmania. A two-way ANOVA confirmed a significant effect of location (*p* < 0.001), but no effect of virus type (*p* = 0.35) and no interaction (*p* = 0.68), indicating that the location effect was similar for both viruses.

### Potential parasitic viruses

A putative full-length 14,902 nucleotide genome of one novel virus was revealed within the *Mononegavirales* order at an abundance of 423 RPM (Figure 9). This virus was related to an invertebrate-infecting virus with 28.72% amino acid similarity (DAZ89743), isolated from an assumed-healthy predatory insect *Gryllotalpa bifratrilecta*^49^, so is unlikely to cause disease in the lamprey, and is likely contamination from the sample collection, either from eukaryotic parasites, diet or even from river-water itself. Geotria australis-associated mononega-like virus did not fall clearly within a known family; therefore, it is classified here as unknown *Mononegavirales*. The genome was annotated using a related virus from the *Artoviridae*, *Brine shrimp artovirus 1*, as a comparison.

**Figure 9.**
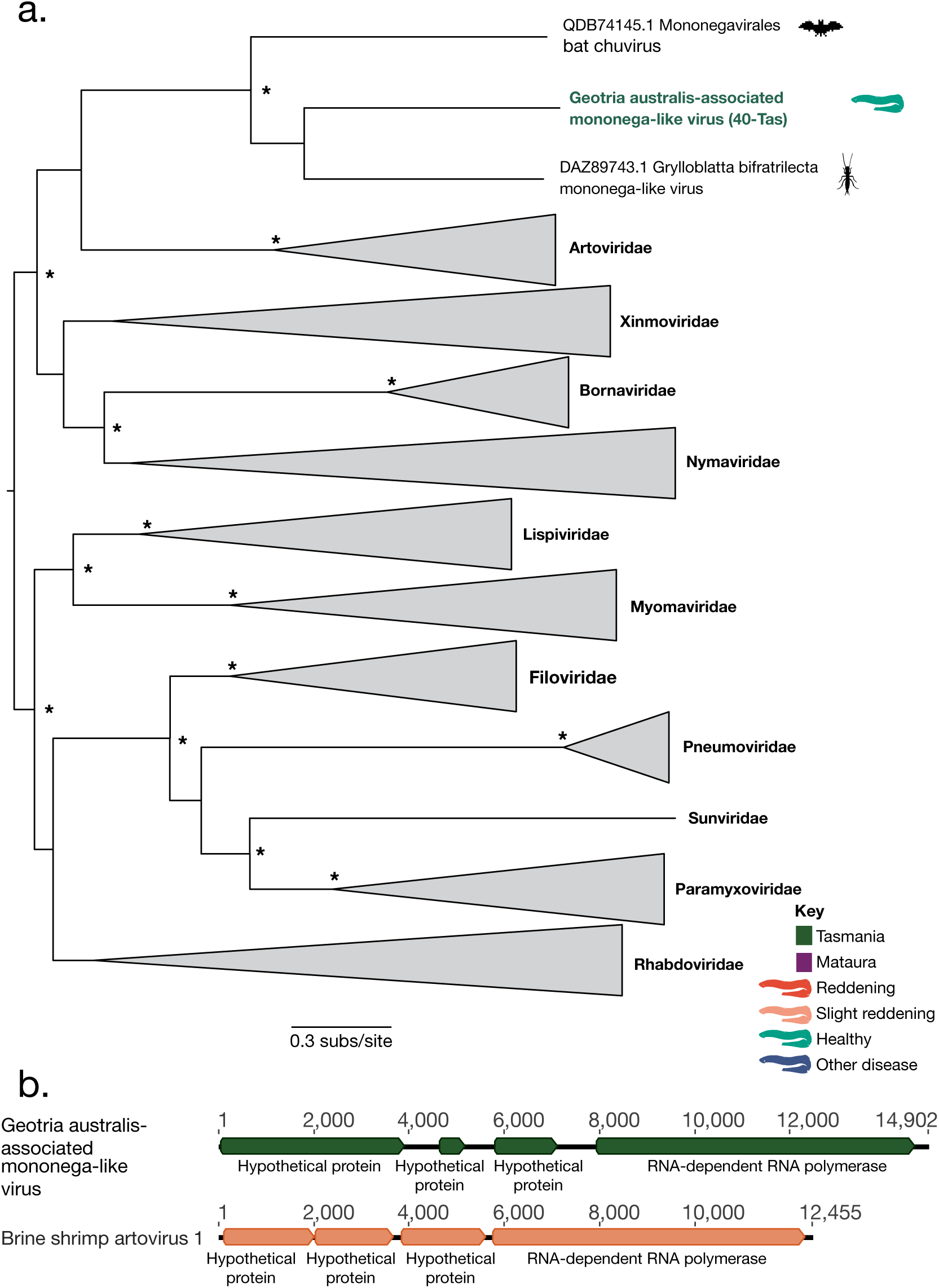
(**a**) Maximum likelihood phylogenetic tree of the Mononegavirales order. An asterisk (*) indicates ultra-fast bootstrap values of ≥95%. Some clades have been collapsed, and trees are mid-point rooted for clarity only. Branch lengths are scaled to the number of amino acid substitutions per site. (**b**) Putative full-length annotated genome of the novel virus compared to a related Brine shrimp artovirus 1 (NCBI accession: OL472418).

## Discussion

This study provides the most comprehensive characterisation to date of the virome associated with lamprey reddening syndrome (LRS) and substantially expands current knowledge of viral diversity in pouched lamprey, *Geotria australis*. Through metatranscriptomic sequencing we identified eight lamprey-associated viruses, seven of which represent novel species. These findings reveal a previously unrecognised diversity of viruses in one of the most evolutionarily ancient vertebrate lineages and provide new insight into the ecological and evolutionary context of LRS.

Lamprey exhibiting reddening harboured significantly higher overall viral abundances, consistent with infection playing a role in disease presentation^50,51^. We detected two highly divergent coronaviruses at high abundance. When analysed separately, *Kanakana letovirus* showed a significant association with disease, whereas Geotria australis coronavirus displayed weaker patterns. These findings are consistent with a scenario in which viral infection precedes or facilitates secondary opportunistic infection by bacteria or oomycetes, ultimately driving tissue pathology. Similar viral-bacterial interactions are well documented in aquatic vertebrates, where polymicrobial dynamics complicate attribution of causation^52,53^.

The recovery of two deeply divergent coronaviruses within a single host species is notable. Geotria australis coronavirus possessed an unusually large, non-segmented genome, while *Kanakana letovirus* had a bi-segmented genome structure, a rare feature within the *Coronaviridae*. The bi-segmented architecture of Kanakana letovirus, demonstrated here for the first time, may enable reassortment and flexible evolutionary dynamics analogous to those seen in other segmented RNA viruses^54–56^. The structural and phylogenetic divergence between these viruses suggests independent host-jump events into lamprey rather than virus-host co-divergence. Their coexistence highlights the evolutionary complexity of the lamprey virome and underscores the deep, largely unexplored history of coronaviruses in aquatic vertebrates.

Beyond coronaviruses, we identified additional highly divergent astroviruses, a picornavirus and an orthomyxovirus, most of which were detected at low abundance and without clear disease association. The presence of diverse, apparently asymptomatic viruses mirrors patterns observed in other fishes and reinforces the need to distinguish infection from disease. The extreme divergence of several viral sequences, sometimes sharing only ∼30% amino acid identity with known relatives, also indicates that considerable viral diversity likely remains undetected in lamprey and other ancient vertebrates. Improvements in remote homology detection and structure-informed approaches may further enhance discovery sensitivity in such evolutionarily distinct hosts^57^.

Geographic comparisons revealed significant differences in virome composition between New Zealand and Australian lamprey, although several viral species were shared between regions, consistent with population connectivity^13^. The apparent predominance of LRS in New Zealand, then, likely reflects ecological or environmental factors rather than strict geographic isolation. Congregation during upstream migration, environmental stressors to pollution or local microbial reservoirs may influence infection dynamics and tipping points for disease expression. The elevated virome burden observed in reddened individuals suggests a threshold in which cumulative viral load overwhelms host defences.

Although sampling location influenced virome composition, interpretation of health-associated patterns is complicated by limited numbers of truly healthy individuals from New Zealand, resulting in potential confounding between location and disease status. However, previous population genomic data indicate that New Zealand and Australian lamprey comprise a single population^13^, and several viral species were shared across regions. Together with the observation that disease associations were driven primarily by differences in viral abundance rather than presence/absence, these results suggest that geographic variation alone is unlikely to explain the patterns observed.

It must be noted that metatranscriptomic detection alone cannot establish pathogenicity, and the presence of a microorganism does not equate to disease^58–62^. Nevertheless, the strong association between LRS and increased coronavirus abundance narrows the list of plausible causative agents. Future work should prioritise targeted assays across larger sample sets and longitudinal sampling to clarify the temporal order of infections. Non-lethal and environmental sampling approaches will be particularly important given the protected status and cultural significance of lamprey^63–64^.

In summary, our findings suggest that LRS is best understood as a polymicrobial syndrome in which coronavirus infection (i.e. *Kanakana letovirus*) may be coupled with host stress from environmental causes or opportunistic bacterial and oomycete infection. While causation remains to be conclusively demonstrated, genomics-informed approaches have resolved longstanding uncertainty surrounding LRS and provided a focused set of viral candidates for future investigation. More broadly, this study expands the known diversity and genomic architecture of viruses infecting ancient vertebrates and illustrates the power of metatranscriptomics to illuminate complex disease processes in threatened wildlife species.

## Data availability

Virus sequences are available on NCBI’s GenBank database under accessions [pending]. The raw reads from lamprey sampled in Australia are available on the SRA under BioProject [pending] while those sampled from New Zealand are available on the Aotearoa Genomics Data Repository [pending].

## Acknowledgements

We thank Jerusha Bennett and Andy Barnes for their assistance with sampling.

## References

1. Costello, M. J. Exceptional endemicity of Aotearoa New Zealand biota shows how taxa dispersal traits, but not phylogeny, correlate with global species richness. J R Soc NZ 54, 144–159 (2024).

2. Grimwood, R. M. et al. From islands to infectomes: host-specific viral diversity among birds across remote islands. BMC Ecol Evol 24, 84 (2024).

3. Hurst, M. R. H., O’Callaghan Maureen, Glare Travis R & and Jackson, T. A. Serratia spp. bacteria evolved in Aotearoa-New Zealand for infection of endemic scarab beetles. N Z J Zool 52, 121–143 (2025).

4. French, R. K. et al. Host phylogeny shapes viral transmission networks in an island ecosystem. Nat. Ecol. Evol. 7, 1834–1843 (2023).

5. Haji, D. et al. Lack of host phylogenetic structure in the gut bacterial communities of New Zealand cicadas and their interspecific hybrids. Sci. Rep. 12, 20559 (2022).

6. Waite, D. W., Deines, P. & Taylor, M. W. Gut Microbiome of the Critically Endangered New Zealand Parrot, the Kakapo (Strigops habroptilus). PloS One 7, e35803 (2012).

7. Waller, S. J. et al. The radiation of New Zealand’s skinks and geckos is associated with distinct viromes. BMC Ecol. Evol. 24, 81 (2024).

8. Brownstein, C. D. & Near, T. J. Phylogenetics and the Cenozoic radiation of lampreys. Curr Biol 33, 397–404.e3 (2023).

9. Gess, R. W., Coates, M. I. & Rubidge, B. S. A lamprey from the Devonian period of South Africa. Nature 443, 981–984 (2006).

10. Miller, A. K. et al. The Southern Hemisphere lampreys (Geotriidae and Mordaciidae). Rev Fish Biol Fish 31, 201–232 (2021).

11. Brosnahan, C. L., Pande, A., Keeling, S. E., van Andel, M. & Jones, J. B. Lamprey (*Geotria australis*; Agnatha) reddening syndrome in Southland rivers, New Zealand 2011–2013: laboratory findings and epidemiology, including the incidental detection of an atypical Aeromonas salmonicida. N Z J Mar Freshw. Res 53, 416–436 (2019).

12. Hilliard, R. W., Pass, D. A. & Potter, I. C. Haemorrhagic Septicaemia in the Lamprey *Geotria australis* Gray. Acta Zool. 60, 115–121 (1979).

13. Miller, A. K. et al. Population Genomics of New Zealand Pouched Lamprey (kanakana; piharau; G*eotria australis*). J Hered 113, (2022).

14. Costa, V. A. et al. Optimising Outbreak Investigation, Surveillance and Discovery of Pathogens in Aquaculture Using Unbiased Metatranscriptomics. Rev. Aquac. 17, e13002 (2025).

15. Doxey, A. C. et al. Metatranscriptomic profiling reveals pathogen and host response signatures of pediatric acute sinusitis and upper respiratory infection. Genome Med 17, 22 (2025).

16. Huang, X. et al. A total infectome approach to understand the etiology of infectious disease in pigs. Microbiome 10, 73 (2022).

17. Robson, M., Chooi, K. M., Blouin, A. G., Knight, S. & MacDiarmid, R. M. A National Catalogue of Viruses Associated with Indigenous Species Reveals High-Throughput Sequencing as a Driver of Indigenous Virus Discovery. Viruses 14, 2477 (2022).

18. Duffy, S., Shackelton, L. A. & Holmes, E. C. Rates of evolutionary change in viruses: patterns and determinants. Nat Rev Genet 9, 267–276 (2008).

19. Landry, M. L. & Owen, M. Failure to Detect Influenza A H1N1 Highlights the Need for Multiple Gene Targets in Influenza Molecular Tests. J Clin Microbiol 61, e0044823 (2023).

20. Osório, N. S. & Correia-Neves, M. Implication of SARS-CoV-2 evolution in the sensitivity of RT-qPCR diagnostic assays. Lancet Infect Dis 21, 166–167 (2021).

21. Taylor, J. T. et al. A Metagenomic Investigation into Apteryx rowi Dermatosis Identifies Multiple Novel Viruses and a Highly Abundant Nematode. J Wildl Dis (2025).

22. Wierenga, J. R. et al. A novel gyrovirus is abundant in yellow-eyed penguin (Megadyptes antipodes) chicks with a fatal respiratory disease. Virology 579, 75–83 (2023).

23. French, R. K. et al. Evidence for a Role of Extraintestinal Pathogenic Escherichia coli, Enterococcus faecalis and Streptococcus gallolyticus in the Aetiology of Exudative Cloacitis in the Critically Endangered Kākāpō (Strigops habroptilus). Mol Ecol 0, e17761 (2025).

24. Fredericks, D. N. & Relman, D. A. Sequence-based identification of microbial pathogens: a reconsideration of Koch’s postulates. Clin Microbiol Rev 9, 18–33 (1996).

25. Byrd, A. L. & Segre, J. A. Adapting Koch’s postulates. Sci. AAAS 351, 224–226 (2016).

26. Ellison, A. R. et al. Comparative transcriptomics reveal conserved impacts of rearing density on immune response of two important aquaculture species. Fish Shellfish Immunol 104, 192–201 (2020).

27. Lv, B.-M., Quan, Y. & Zhang, H.-Y. Causal Inference in Microbiome Medicine: Principles and Applications. Trends Microbiol 29, 736–746 (2021).

28. Hutson, K. S., Davidson, I. C., Bennett, J., Poulin, R. & Cahill, P. L. Assigning cause for emerging diseases of aquatic organisms. Trends Microbiol 31, 681–691 (2023).

29. Lane, H. S., Brosnahan, C. L. & Poulin, R. Aquatic disease in New Zealand: synthesis and future directions. N. Z. J. Mar. Freshw. Res. 56, 1–42 (2022).

30. Ruben, M. O., Akinsanola, A. B., Okon, M. E., Shitu, T. & Jagunna. Emerging challenges in aquaculture: Current perspectives and human health implications. Vet World 18, 15–28 (2025).

31. Bolger, A. M., Lohse, M. & Usadel, B. Trimmomatic: a flexible trimmer for Illumina sequence data. Bioinformatics 30, 2114–2120 (2014).

32. Andrews, S. FastQC: a quality control tool for high throughput sequence data. (2010).

33. Li, D., Liu, C.-M., Luo, R., Sadakane, K. & Lam, T.-W. MEGAHIT: an ultra-fast single-node solution for large and complex metagenomics assembly via succinct de Bruijn graph. Bioinformatics 31, 1674–1676 (2015).

34. Langmead, B. & Salzberg, S. L. Fast gapped-read alignment with Bowtie 2. Nat. Methods 9, 357–359 (2012).

35. Oksanen, J. et al. Vegan: Community Ecology Package. R Package Version 22-1 **2**, 1–2 (2015).

36. Derek H. Ogle, Jason C. Doll, A. Powell Wheeler & Alexis Dinno. FSA: Simple Fisheries Stock Assessment Methods. 10.32614/CRAN.package.FSA(2025) doi:10.32614/CRAN.package.FSA.

37. Hadley Wickham. Ggplot2: Elegant Graphics for Data Analysis. (Springer-Verlag New York, 2016).

38. Katoh K, Misawa K, Kuma K, Miyata T. MAFFT: a novel method for rapid multiple sequence alignment based on fast Fourier transform. Nucleic Acids Res. 30(14):3059–66 (2002).

39. Capella-Gutiérrez S, Silla-Martínez JM, Gabaldón T. trimAl: a tool for automated alignment trimming in large-scale phylogenetic analyses. Bioinformatics, 25(15):1972–3 (2009).

40. Nguyen LT, Schmidt HA, von Haeseler A, Minh BQ. IQ-TREE: a fast and effective stochastic algorithm for estimating maximum-likelihood phylogenies. Mol Biol Evol, 32(1):268–74 (2015).

41. Söding, J., Biegert, A. & Lupas, A. N. The HHpred interactive server for protein homology detection and structure prediction. Nucleic Acids Res 33, W244–8 (2005).

42. Neuman, B. W. et al. Giant RNA genomes: Roles of host, translation elongation, genome architecture, and proteome in nidoviruses. Proc. Natl. Acad. Sci. 122, e2413675122 (2025).

43. Petrone, M. E., et al. Evidence for an ancient aquatic origin of the RNA viral order Articulavirales. Proc. Natl. Acad. Sci. - PNAS 120, 1-e2310529120 (2023).

44. Falchieri M, Coward VJ, Reid SM, Lewis T, Banyard AC. Infectious bronchitis virus: an overview of the “chicken coronavirus”. J Med Microbiol, 73(5):001828 (2024).

45. Miller, A. K. et al. Slippery when wet: cross-species transmission of divergent coronaviruses in bony and jawless fish and the evolutionary history of the *Coronaviridae*. Virus Evol 7, (2021).

46. Lauber C, Zhang X, Vaas J, Klingler F, Mutz P, Dubin A, Pietschmann T, Roth O, Neuman BW, Gorbalenya AE, Bartenschlager R, Seitz S. Deep mining of the Sequence Read Archive reveals major genetic innovations in coronaviruses and other nidoviruses of aquatic vertebrates. PLoS Pathog, 20(4):e1012163 (2024).

47. Šimičić P, Židovec-Lepej S. A Glimpse on the Evolution of RNA Viruses: Implications and Lessons from SARS-CoV-2. Viruses, 15(1):1 (2022).

48. Gorbalenya, A. E., Enjuanes, L., Ziebuhr, J. & Snijder, E. J. Nidovirales: evolving the largest RNA virus genome. Virus Res 117, 17–37 (2006).

49. Wu H, Pang R, Cheng T, Xue L, Zeng H, Lei T, Chen M, Wu S, Ding Y, Zhang J, Shi M, Wu Q. Abundant and diverse RNA viruses in insects revealed by RNA-Seq analysis: Ecological and evolutionary implications. mSystems, 5(4):e00039–20 (2020).

50. Fajnzylber, J. et al. SARS-CoV-2 viral load is associated with increased disease severity and mortality. Nat Commun 11, 5493 (2020).

51. Kim, M.-J., Kim, J.-O., Jang, G.-I., Kwon, M.-G. & Kim, K.-I. Evaluation of the horizontal transmission of white spot syndrome virus for whiteleg shrimp (Litopenaeus vannamei) based on the disease severity grade and viral shedding rate. Animals 13, 1676 (2023).

52. Okon, E. M., Okocha, R. C., Taiwo, A. B., Michael, F. B. & Bolanle, A. M. Dynamics of co-infection in fish: A review of pathogen-host interaction and clinical outcome. Fish Shellfish Immunol Rep 4, 100096 (2023).

53. Basri, L. et al. Co-Infections of tilapia lake virus, Aeromonas hydrophila and Streptococcus agalactiae in farmed red hybrid tilapia. Animals 10, 2141 (2020).

54. Pérez-Losada, M., Arenas, M., Galán, J. C., Palero, F. & González-Candelas, F. Recombination in viruses: mechanisms, methods of study, and evolutionary consequences. Infect Genet Evol 30, 296–307 (2015).

55. Kim, H., Webster, R. G. & Webby, R. J. Influenza Virus: Dealing with a Drifting and Shifting Pathogen. Viral Immunol 31, 174–183 (2018).

56. Petrova, V. N. & Russell, C. A. The evolution of seasonal influenza viruses. Nat. Rev. Microbiol. 16, 47–60 (2018).

57. Wu, F., Janvier, P. & Zhang, C. The rise of predation in Jurassic lampreys. Nat Commun 14, 6652 (2023).

58. Grimwood, R. M., Holmes, E. C. & Geoghegan, J. L. A Novel Rubi-Like Virus in the Pacific Electric Ray (Tetronarce californica) Reveals the Complex Evolutionary History of the Matonaviridae. Viruses 13, 585 (2021).

59. Costa, V. A. et al. Host adaptive radiation is associated with rapid virus diversification and cross-species transmission in African cichlid fishes. Curr. Biol. 34, 1247–1257.e3 (2024).

60. Geoghegan, J. L. et al. Hidden diversity and evolution of viruses in market fish. Virus Evol 4, (2018).

61. Costa, V. A. & Holmes, E. C. Diversity, evolution, and emergence of fish viruses. J Virol 98, e00118–24 (2024).

62. Grimwood, R. M. et al. Viromes of Antarctic fish resemble the diversity found at lower latitudes. Virus Evol 10, (2024).

63. Williams, E. et al. Understanding taonga freshwater fish populations in Aotearoa–New Zealand. NIWA Client Rep. 2017326HN, 1–228 (2017).

64. Todd, P. R. A status report on the New Zealand lamprey. N.Z. Freshw. Fish. Misc. Rep. 11, 1–32 (1992).

