## Supplementary Table 1 for "Divergent coronaviruses discovered in the virome of lamprey with reddening syndrome"

**Supplementary Table 1**. Meta data of the 28 pooled RNA libraries

, including sampling location, health status, and concentration, including 260/280 ratio measured on a Nanodrop Spectrophotometer. A tick indicates the tissues pooled into the library.

| Library ID | Sampling location | Health status | Concentration (ng/µL) | 260/280 | No. tissues in pool | Skin | Gill | Liver | Muscle | Kidney |
| --- | --- | --- | --- | --- | --- | --- | --- | --- | --- | --- |
| 2024-35-Tas-D | Tasmania | Diseased | 146 | 2.09 | 5 | ✓ | ✓ | ✓ | ✓ | ✓ |
| 2024-59-Tas-D | Tasmania | Diseased | 127.4 | 2.05 | 5 | ✓ | ✓ | ✓ | ✓ | ✓ |
| 2024-1-Gore-H | Mataura | Healthy | 175.3 | 2.07 | 4 | ✓ | ✓ | ✓ |  | ✓ |
| 2024-2-Gore-H | Mataura | Healthy | 91 | 2.11 | 4 | ✓ | ✓ | ✓ |  | ✓ |
| 2024-31-Tas-H | Tasmania | Healthy | 157.1 | 2.08 | 4 | ✓ | ✓ | ✓ |  | ✓ |
| 2024-32-Tas-H | Tasmania | Healthy | 67.6 | 2.09 | 5 | ✓ | ✓ | ✓ | ✓ | ✓ |
| 2024-33-Tas-H | Tasmania | Healthy | 229.8 | 2.09 | 4 | ✓ |  | ✓ | ✓ | ✓ |
| 2024-34-Tas-H | Tasmania | Healthy | 117.2 | 2.09 | 5 | ✓ | ✓ | ✓ | ✓ | ✓ |
| 2024-36-Tas-H | Tasmania | Healthy | 150.3 | 2.09 | 5 | ✓ | ✓ | ✓ | ✓ | ✓ |
| 2024-37-Tas-H | Tasmania | Healthy | 104.6 | 2.06 | 5 | ✓ | ✓ | ✓ | ✓ | ✓ |
| 2024-38-Tas-H | Tasmania | Healthy | 121.7 | 2.05 | 5 | ✓ | ✓ | ✓ | ✓ | ✓ |
| 2024-39-Tas-H | Tasmania | Healthy | 320.5 | 2.09 | 5 | ✓ | ✓ | ✓ | ✓ | ✓ |
| 2024-40-Tas-H | Tasmania | Healthy | 223.4 | 2.08 | 5 | ✓ | ✓ | ✓ | ✓ | ✓ |
| 2024-17-Gore-H | Mataura | Healthy with slight reddening | 576.6 | 2.04 | 5 | ✓ | ✓ | ✓ | ✓ | ✓ |
| 2024-18-Gore-H | Mataura | Healthy with slight reddening | 690.5 | 2.06 | 5 | ✓ | ✓ | ✓ | ✓ | ✓ |
| 2024-19-Gore-H | Mataura | Healthy with slight reddening | 202.9 | 2.1 | 5 | ✓ | ✓ | ✓ | ✓ | ✓ |
| 2024-20-Gore-H | Mataura | Healthy with slight reddening | 330.4 | 2.09 | 5 | ✓ | ✓ | ✓ | ✓ | ✓ |
| 2024-12-Gore-D | Mataura | Reddening | 160.6 | 2.09 | 5 | ✓ | ✓ | ✓ | ✓ | ✓ |
| 2024-13-Gore-D | Mataura | Reddening | 150.2 | 2.08 | 5 | ✓ | ✓ | ✓ | ✓ | ✓ |
| 2024-14-Gore-D | Mataura | Reddening | 326 | 2.09 | 5 | ✓ | ✓ | ✓ | ✓ | ✓ |
| 2024-15-Gore-D | Mataura | Reddening | 312.5 | 2.09 | 4 | ✓ | ✓ | ✓ | ✓ |  |
| 2024-16-Gore-D | Mataura | Healthy with slight reddening | 423 | 2.1 | 4 | ✓ | ✓ | ✓ | ✓ |  |
| 2024-3-Gore-D | Mataura | Reddening | 106.2 | 2.08 | 4 | ✓ | ✓ | ✓ |  | ✓ |
| 2024-4-Gore-D | Mataura | Reddening | 218 | 2.1 | 3 | ✓ | ✓ | ✓ |  |  |
| 2024-5-Gore-D | Mataura | Reddening | 168 | 2.07 | 5 | ✓ | ✓ | ✓ | ✓ | ✓ |
| 2024-6-Gore-D | Mataura | Reddening | 81.9 | 2.08 | 5 | ✓ | ✓ | ✓ | ✓ | ✓ |
| 2024-7-Gore-D | Mataura | Reddening | 2.09 | 2.1 | 5 | ✓ | ✓ | ✓ | ✓ | ✓ |
| 2024-8-Gore-D | Mataura | Reddening | 316 | 2.1 | 5 | ✓ | ✓ | ✓ | ✓ | ✓ |
