## Supplementary Table 2 for "Divergent coronaviruses discovered in the virome of lamprey with reddening syndrome"

**Supplementary Table 2**. Fish-infecting viruses identified in this study and their closest known genetic relative from a BLASTp search. Details include proposed virus names following collaboration with mana whenua.

| Library | Disease status | | Virus name | Viral abundance (RPM) | ORF (AA) | Full genome (nt) | Family | Location | Top BLASTp hit name | E value | Amino acid percentage identity | Top hit accession number |
| --- | --- | --- | --- | --- | --- | --- | --- | --- | --- | --- | --- | --- |
| 2024-38-Tas-H | Healthy | | Geotria australis piscistrovirus 1 | 0.294273227 | 212 | NA | Astroviridae | Tasmania | *Jingmen bat astrovirus 1* | 4.00E-17 | 30.61% | WOK58275.1 |
| 2024-31-Tas-H | Healthy | | Geotria australis piscistrovirus 1 | 0.131410839 | 130 | NA |  | Tasmania | *Wenling gobies fish astro-like virus* | 1.00E-10 | 29.46% | AVM87598.1 |
| 2024-12-Gore-D | Reddening | | Geotria australis piscistrovirus 2 | 0.337705306 | 178 | NA | Astroviridae | Gore | *Jingmen bat astrovirus 1* | 3.00E-32 | 45.62% | WOK58275.1 |
| 2024-14-Gore-D | Reddening | | Geotria australis piscistrovirus 3 | 0.239845268 | 180 | NA | Astroviridae | Gore | *Jingmen bat astrovirus 1* | 3.00E-33 | 45.78% | WOK58275.1 |
| 2024-14-Gore-D | Reddening | | Geotria australis piscistrovirus 4 | 0.125633236 | 148 | NA | Astroviridae | Gore | *Porure astrovirus 1* | 5.00E-08 | 31.03% | UNJ19229.1 |
| 2024-12-Gore-D | Reddening | | Geotria australis piscistrovirus 4 | 0.168852653 | 144 | NA | Astroviridae | Gore | *Porure astrovirus 1* | 1.00E-20 | 40.17% | UNJ19228.1 |
| 2024-16-Gore-D | Healthy with slight reddening | Geotria australis piscistrovirus 4 | | 0.273616946 | 233 | NA | Astroviridae | Gore | *Xenotilapia nasus astrovirus 2* | 5.00E-16 | 27.83% | WLN26242.1 |
| 2024-19-Gore-H | Healthy with slight reddening | Geotria australis piscistrovirus 4 | | 0.267293715 | 221 | NA | Astroviridae | Gore | *Xenotilapia nasus astrovirus 2* | 5.00E-15 | 26.87% | WLN26242.1 |
| 2024-38-Tas-H | Healthy | | Geotria australis piscistrovirus 4 | 0.101473527 | 132 | NA | Astroviridae | Tasmania | *Limnochromis auritus astrovirus* | 3.00E-23 | 39.39% | WLN26244.1 |
| 2024-34-Tas-H | Healthy | | Geotria australis piscistrovirus 4 | 0.200656616 | 126 | NA | Astroviridae | Tasmania | *Limnochromis auritus astrovirus* | 4.00E-23 | 43.09% | WLN26244.1 |
| 2024-31-Tas-H | Healthy | | Geotria australis piscistrovirus 4 | 0.058404817 | 108 | NA | Astroviridae | Tasmania | *Xenotilapia nasus astrovirus 2* | 2.00E-06 | 29.91% | WLN26242.1 |
| 2024-37-Tas-H | Healthy | | Geotria australis piscistrovirus 4 | 0.345126758 | 192 | NA | Astroviridae | Tasmania | *Xenotilapia nasus astrovirus 2* | 9.00E-18 | 29.38% | WLN26242.1 |
| 2024-16-Gore-D | Healthy with slight reddening | | Geotria australis influenza-like virus | 0.517917791 | 488 | NA | Orthomyxoviridae | Gore | *Sturgeon influenza-like virus* | 3.00E-71 | 35.83% | DBA45437.1 |
| 2024-31-Tas-H | Healthy | | Geotria australis picormavirus | 1.628034277 | 1029 | NA | Picornavirales | Tasmania | *Picornavirales sp.* | 1.00E-75 | 32.34% | QJI53587.1 |
| 2024-59-Tas-H | Unknown disease | | Geotria australis coronavirus | 2.22 | 6211 | 35,232 | Coronaviridae | Tasmania | *Bat coronavirus* | 2e-32 | 26.32% | XYH51711.1 |
| 2024-12-Gore-D | Reddening | | Geotria australis coronavirus | 54.04 | 5374 | 35,576 | Coronaviridae | Gore | *Alphacoronavirus sp.* | 3e-105 | 25.47% | QGX41962.1 |
| 2024-07-Gore-D | Reddening | | Geotria australis coronavirus | 50.09 | 6237 | 35,609 | Coronaviridae | Gore | *Bat coronavirus* | 7e-63 | 28,39% | XYH51711.1 |
| 2024-05-Gore-D | Reddening | | Geotria australis coronavirus | 1973.9 | 6237 | 35,594 | Coronaviridae | Gore | *Alphacoronavirus sp.* | 2e-118 | 26.05% | QGX41962.1 |
| 2024-20-Gore-H | Healthy with slight reddening | Geotria australis coronavirus | | 725.13 | 6237 | 35,586 | Coronaviridae | Gore | *Alphacoronavirus sp.* | 2e-118 | 26.65% | QGX41962.1 |
| 2024-02-Gore-H | Healthy | | Geotria australis coronavirus | 584.08 | 6237 | 35,628 | Coronaviridae | Gore | *Alphacoronavirus sp.* | 1e-118 | 26.59% | QGX41962.1 |
| 2024-14-Gore-D | Reddening | | Geotria australis coronavirus | 28.18 | 5343 | 35,558 | Coronaviridae | Gore | *Swan goose coronavirus* | 1e-47 | 22.50% | WKW84054.1 |
| 2024-08-Gore-D | Reddening | | *Kanakana letovirus* | 10.99 | 4431 | 31,992 | Coronaviridae | Gore | *Kanakana letovirus* | 0.0 | 94.14% | UBF41764.1 |
| 2024-04-Gore-D | Reddening | | *Kanakana letovirus* | 23.89 | 4228 | 31,849 | Coronaviridae | Gore | *Kanakana letovirus* | 0.0 | 98.12% | UBF41764.1 |
| 2024-01-Gore-H | Healthy | | *Kanakana letovirus* | 91.1 | 4228 | 31,850 | Coronaviridae | Gore | *Kanakana letovirus* | 0.0 | 99.26% | UBF41764.1 |
| 2024-06-Gore-D | Reddening | | *Kanakana letovirus* | 40.17 | 4264 | 31,918 | Coronaviridae | Gore | *Kanakana letovirus* | 0.0 | 95.63% | UBF41764.1 |
| 2024-03-Gore-D | Reddening | | *Kanakana letovirus* | 78.42 | 4228 | 31,850 | Coronaviridae | Gore | *Kanakana letovirus* | 0.0 | 99.26% | UBF41764.1 |
| 2024-12-Gore-D | Reddening | | *Kanakana letovirus* | 187.76 | 4228 | 31,879 | Coronaviridae | Gore | *Kanakana letovirus* | 0.0 | 99.16% | UBF41764.1 |
| 2024-07-Gore-D | Reddening | | *Kanakana letovirus* | 14.22 | 4204 | 31,182 | Coronaviridae | Gore | *Kanakana letovirus* | 0.0 | 84.17% | UBF41764.1 |
